# Neuronal Responses in Posterior Parietal Cortex during Learning of Implied Serial Order

**DOI:** 10.1101/689133

**Authors:** Fabian Munoz, Greg Jensen, Benjamin C. Kennedy, Yelda Alkan, Herbert S. Terrace, Vincent P. Ferrera

## Abstract

Monkeys are able to learn the implied ordering of pairs of images drawn from an ordered set, without ever seeing all of the images simultaneously and without explicit spatial or temporal cues. The learning of implied order differs from learning of explicit visual or motor sequences. We recorded the activity of parietal neurons in rhesus macaques while they learned 7-item TI lists when only 2 items were presented on each trial. Behavior and ensemble neuronal activity were significantly influenced by the ordinal relationship of the stimulus pairs, specifically symbolic distance (the difference in rank) and joint ranks (the sum of the ranks). Symbolic distance strongly predicted decision accuracy, and learning was consistently faster as symbolic distance increased. An effect of joint rank on performance was also found nested within the symbolic distance effect. Across the population of neurons, there was significant modulation of firing correlated with the relative ranks of the two stimuli presented on each trial. Neurons exhibited selectivity for stimulus rank during learning, but not before or after. The observed behavior during learning is best explained by a virtual workspace model, not by associative or reward mechanisms. The neural data support a role for posterior parietal cortex in representing several variables that contribute to serial learning, particularly information about the ordinal ranks of the stimuli presented during a given trial. Thus, parietal cortex appears to belong to a neural substrate for learning and representing abstract relationships in a virtual workspace.

---

Transitive inference (TI) refers to judgments that if A>B and B>C, then one can infer that A>C by the transitive property of ordinal rank (Halford et al., 2010). Rank efficiently encodes list order or hierarchy, as it requires only one value for each item (which scales linearly with list length), rather than encoding a separate relationship for each combination of items (which scales exponentially). A wide variety of species can learn ordered lists (Vasconcelos, 2008; Jensen, 2017). Such hierarchical knowledge is considered important in a variety of natural contexts (Moses et al., 2006; Haun et al., 2010; Gazes et al., 2013).

Several lines of evidence suggest that abstract cognitive mechanisms give rise to TI abilities (Frank et al., 2006; Kumaran and Ludwig, 2013), distinguishing them from the learning of temporal or motor sequences. The strongest of these is the presence of a symbolic distance effect (SDE). Symbolic distance refers to the difference in the ranks of the two items (e.g. adjacent pairs have SD=1), and decision accuracy varies directly with the symbolic distance that separates each pair (Terrace et al., 2003; Terrace, 2012). Recent evidence suggests that purely associative mechanisms and reward signals cannot account for these effects (Jensen et al., 2013; Kumaran, 2013; Jensen et al., 2019). Abstract representations of ordinal rank may be linked to those of spatial position (Roberts and Phelps, 1994; Taffe et al., 2004), with both utilizing a “virtual workspace.” This workspace may be used across modalities, such that spatial and non-spatial information are integrated into a common representation (Gevers et al., 2003; Walsh, 2003; Hubbard et al., 2005). Such a workspace would be a computationally efficient solution to inferring the implied ordering of stimuli, especially when temporal or spatial cues provide no information about that ordering.

Parietal cortex has long been associated with spatial relationships, and parietal lobe damage in humans gives rise to a well-known spatial hemineglect syndrome. In parietal stroke cases (e.g. Gerstmann Syndrome), multimodality spatial and number-related deficits can co-occur (Pia et al., 2009). Parietal dysfunction involving the intraparietal sulcus (IPS) is associated with irregularities in spatial and numerical representation (Wingard et al., 2002; Molko et al., 2003). Posterior parietal cortex (PPC) has been shown to integrate spatial, temporal, and reward information (Roitman et al., 2007). There is also evidence suggesting that the same areas of parietal lobe are responsible for both spatial representation and serial learning (Acuna et al., 2002; Hinton et al., 2010; Smets et al., 2013), including TI (Gould et al., 2009; Prado et al., 2013).

IPS is also implicated in various number representation and manipulation tasks (Nieder et al., 2002; Piazza et al., 2004, 2007; Prado et al., 2010; Roitman et al., 2012). Electrophysiological studies suggest that PPC encodes numerosity (Jacob and Nieder, 2009; Sawamura et al., 2010; Viswanathan and Nieder, 2013). Lateral intraparietal (LIP) neurons exhibit features of a magnitude accumulator (Dehaene and Changeux, 1993; Sawamura et al., 2002; Nieder, 2013) and are implicated in spatial cognition, including spatial attention (Bisley and Goldberg, 2003), judging relative spatial position (Chafee et al., 2007), and probabilistic reasoning (Yang and Shadlen, 2007). LIP also appears to integrate relevant spatial, temporal, numerical, and reward signals guiding behavior (Freedman and Assad, 2009), making this region a candidate for representation of the virtual workspace.

Electrophysiology in non-human primates offers an opportunity to study how the implied ordering of a set of stimuli is learned without confounds of semantic encoding and language use. Because macaques can learn 7-item lists within a few hundred trials, their behavior can be examined with concurrent single-unit electrophysiology. The present study is the first to examine the electrophysiological basis of ordinal position and distance effects during the training of tasks in which the stimulus order is entirely implicit. Evidence that the firing rates of LIP neurons correlate with ordinal position of or symbolic distance between stimuli, would support the encoding of a mental number line in LIP. Our paradigm is therefore uniquely poised to address whether TI is accomplished by using purely associative mechanisms, and, if not, how higher order representations in parietal cortex might be involved.

## MATERIALS & METHODS

### Subjects

Subjects were two male rhesus macaques (H and L), 13-14 years old and weighing 9-10 kg at time of experiment. The research was conducted in accordance with U.S. Department of Health and Human Services (National Institutes of Health) guidelines for the care and use of laboratory animals, and was approved by the Institutional Animal Care and Use Committee at Columbia University and the New York State Psychiatric Institute. Monkeys were prepared for experiments by surgical implantation of a post for head restraint and a recording chamber to give access to cortex. All surgery was performed under general anesthesia (isoflurane 1-3%) and aseptic conditions. Subjects had prior experience performing transitive inference tasks (Jensen et al., 2015).

### Visual Stimuli & Eye Movements

The subjects were seated in an upright primate chair during the experiment and responded to visual stimuli by making saccadic eye movements. Stimuli were generated and controlled by a CRS VSG2/3F video frame buffer. Stimuli were displayed on a CRT (subject H) or LCD (subject L) monitor with a resolution of 1280 x 1024 pixels at 60 Hz. Viewing distance was 60 cm.

Visual stimuli consisted of 140-by-130 pixel color photographs (7°by 8°visual angle) and small squares that served as fixation and eye movement targets. The fixation point was a red square (0.5°visual angle). Eye position was recorded using a monocular scleral search coil (CNC Engineering, Seattle WA) and digitized at a rate of 1 kHz (Judge et al., 1980). Subjects made choices by making eye movements from the central fixation point to target stimuli. Eye velocity was computed offline by convolving eye position with a digital filter. The filter was constructed by taking the first derivative of a temporal Gaussian, *G*(*t*), such that

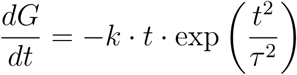

Here, *τ* = 8 msec and *k* is a constant that sets the filter gain to 1.0. This filter does not introduce a time shift between the position input and velocity output, but adds temporal uncertainty to the velocity estimates. Horizontal and vertical eye velocities (*h*′(*t*) and *v*′(*t*), respectively) were combined to estimate radial eye speed *r*′(*t*) using the Pythagorean identity:

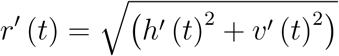

### Behavioral Tasks

Subjects were trained to perform three visual-oculomotor tasks: delayed visually guided saccades, delayed memory guided saccades, and transitive inference.

#### Delayed visual and memory guided saccades

The purpose of this task was to estimate the spatial selectivity (receptive and movement field) of each neuron. Each trial started with a small white spot (0.5°white square) presented in the center of the video display. Once the subject fixated on this spot, a peripheral cue appeared (1.0°white square). The subject was required to hold fixation for 0.75 to 1.25 seconds. Then the central fixation target was turned off, instructing the subject to make a saccade to the peripheral cue. The eccentricity and azimuth of the peripheral cue varied from trial to trial. Targets were presented at 8, 16, or 24 locations. This allowed the experimenter to determine the spatial locations to which each neuron responded most strongly. The stimulus presented at the preferred location is hereafter referred to as falling within the cell’s receptive field. The other stimulus was presented diametrically opposite relative to the center of the screen, and is referred to as falling outside the receptive field. The delay between the peripheral cue onset and the saccade allowed experimenters to distinguish between visual and movement-related neuronal activity. The memory guided saccade task was identical to the visually guided saccade task, except that the peripheral cue was presented only briefly (0.5 sec) and then turned off. During the cue presentation and the subsequent memory delay interval (0.75 to 1.25 sec), the subject continued to fixate on the center fixation target. Then the central fixation target was turned off, instructing the subject to make a saccade to the remembered location of the peripheral cue. After the saccade, the peripheral cue was turned back on to provide feedback about saccade accuracy.

#### Transitive inference

Prior to the beginning of each recording session, a set of 7 photographs never before seen by the subjects was selected from a database of over 2500 images. A different set of images was used for each session. These stimuli were randomly assigned a unique rank, indicated by the letters A-G, to create an ordered list (Fig. 1A). No explicit information about stimulus rank was presented to the subjects. That is, the letter assigned to each stimulus was never displayed, nor was there any information about serial order in the spatial or temporal presentation of the stimuli.

**Figure 1.**
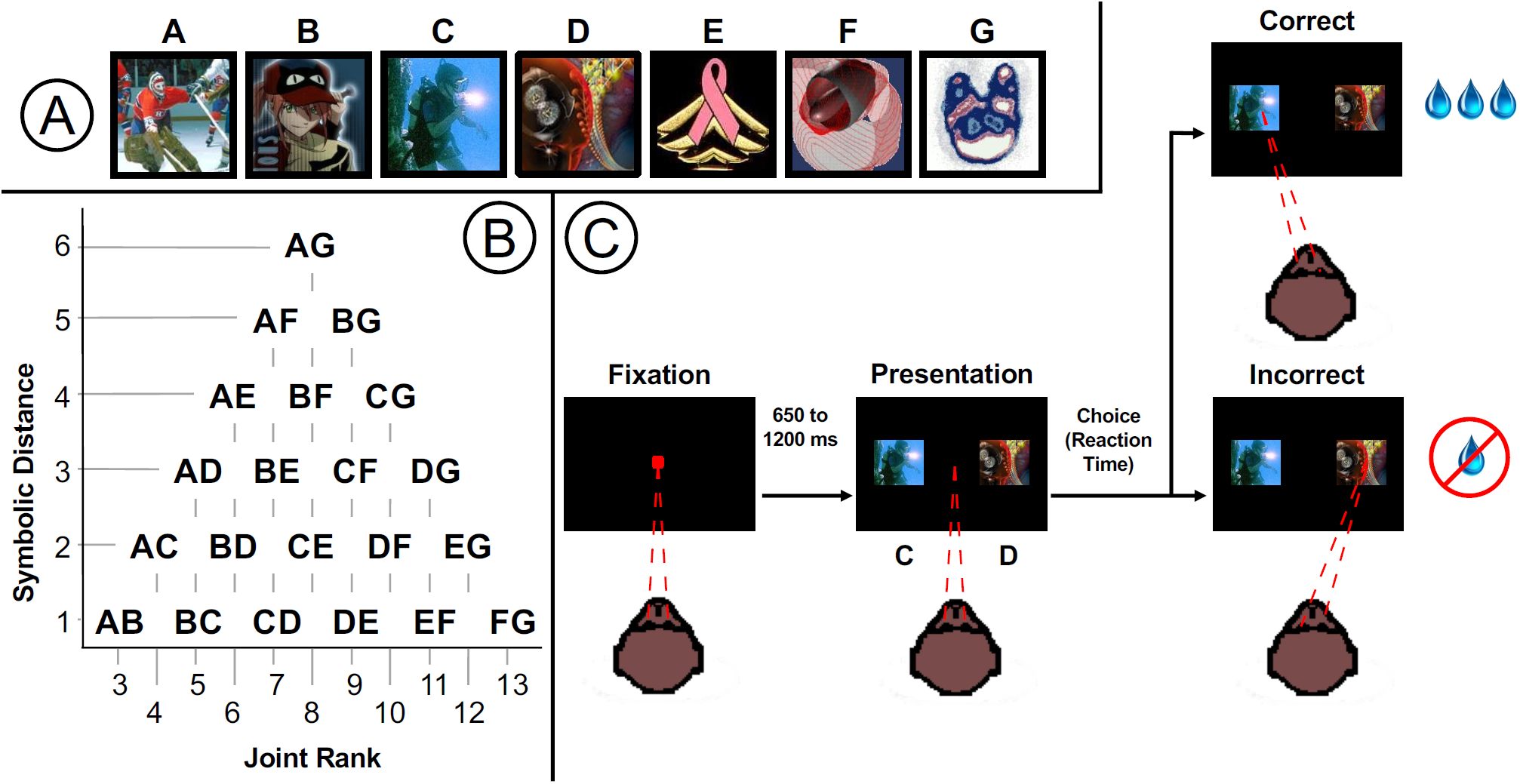
Transitive inference task procedure. **A.** Example of a list of 7 random images assigned ranks A thru G. **B.** All 21 image pairs for a 7-item list, and their respective joint ranks and symbolic distances. **C.** Trials structure and timing.

For 7-item lists, there are 21 possible pairs. In addition to being identified in terms of which stimuli they included, each pair could be described in terms of two metrics that encode information about both stimuli. The “symbolic distance” was calculated by taking the difference between the stimulus ranks (D’Amato and Colombo, 1990). For example, since the pair BC consists of items that are adjacent in the list (i.e. ranks 2 and 3, respectively), their symbolic distance is 1, while the pair AG (ranks 1 and 7) has a symbolic distance of 6. The “joint rank” is the sum of the ranks of the two stimuli presented on each trial, and is orthogonal to symbolic distance (Jensen et al., 2017). BC would have a joint rank of 5 (2+3), while AG would have a joint rank of 8 (1+7). A diagram mapping the symbolic distance and joint rank for all pairs is presented in Figure 1B.

During each trial, the subject first fixated a small target in the center of the screen (Fig. 1C). Following a random delay (0.4 sec to 1.2 sec, positively skewed with a mean of 0.5 sec), the fixation point disappeared and two stimuli appeared equidistant from the center of the screen in opposite directions. To receive a reward, subjects had to select the stimulus with the lower rank by making a saccadic eye movement to the selected stimulus within 1.5 sec of stimulus onset, and fixating for a random interval (0.4 to 0.6 sec, uniformly distributed). When this interval elapsed, auditory feedback was delivered indicating that the response was either correct (high tone, 880 Hz) and would be rewarded, or was incorrect (low tone, 440 Hz). In order to receive the reward on correct trials, subjects had to maintain fixation for another interval (0.35 to 0.65 sec, uniformly distributed), after which the screen went dark and fluid rewards (juice or water drops) were delivered.

The first phase of each session presented single stimuli from the list. A “block” of responses during this phase consisted of presentations of each of the stimuli, both in and out of the cell’s receptive field. Thus, for a 7-item list, each block was 14 trials long, randomly ordered. The first phase lasted 5 blocks (70 trials total.) A saccade to these stimuli always yielded a reward. These presentations were used to determine whether the cells demonstrated selectivity toward any of the individual stimuli prior to training. For the remainder of the session, pairs of stimuli were presented. With positional counterbalancing, this resulted in blocks that were 42 trials in length. In order to receive a reward, subjects had to choose the stimulus with the lower rank. Subjects were presented with all pairs of stimuli, one randomized block at a time, for up to 20 blocks (i.e. 840 trials in total).

If good isolation from the recorded cell was maintained past the end of the all-pairs phase, and the subject remained motivated to work, a new list of photographs was drawn from the database, and the task was repeated (with another 70 trials of single stimuli and up to 20 blocks of pairs). During a single recording session, subjects were capable of learning 1 to 3 lists in this fashion.

### Neuronal Recording

Subjects had plastic recording chambers implanted over the right hemisphere at stereotaxic coordinates (subject H: anterior 5, medial 16; subject L: posterior 6, medial 14). Single-cell activity was recorded with glass-coated tungsten electrodes (Alpha Omega) with impedances between 0.5 and 2 MΩ measured at a frequency of 1 kHz. Raw signals were amplified, digitized, and high-pass filtered. We used FHC APM and Alpha Omega SnR recording systems. Action potential waveforms were identified by a time-amplitude window discriminator (FHC preamplifier) or threshold crossings (Alpha Omega). Action potentials were converted into digital pulses that were sampled by the computer with 0.02 ms temporal resolution. Waveforms were stored for offline spike sorting using WaveSorter (Phillips, 2012) or WaveClus (Quiroga et al., 2004). However, offline spike sorting was necessary only in exceptional cases because recording was mostly restricted to cells that could be clearly isolated online.

Recording sites were anatomically reconstructed using combination of recording grid coordinates, microdrive depth readings, and stereotaxic position of the chamber, and were projected onto the structural MRI using custom MATLAB scripts. Figure 2 plots a reconstruction of recording sites on a template atlas of the anatomical macaque rhesus brain for area classification (Reveley et al., 2017).

**Figure 2.**
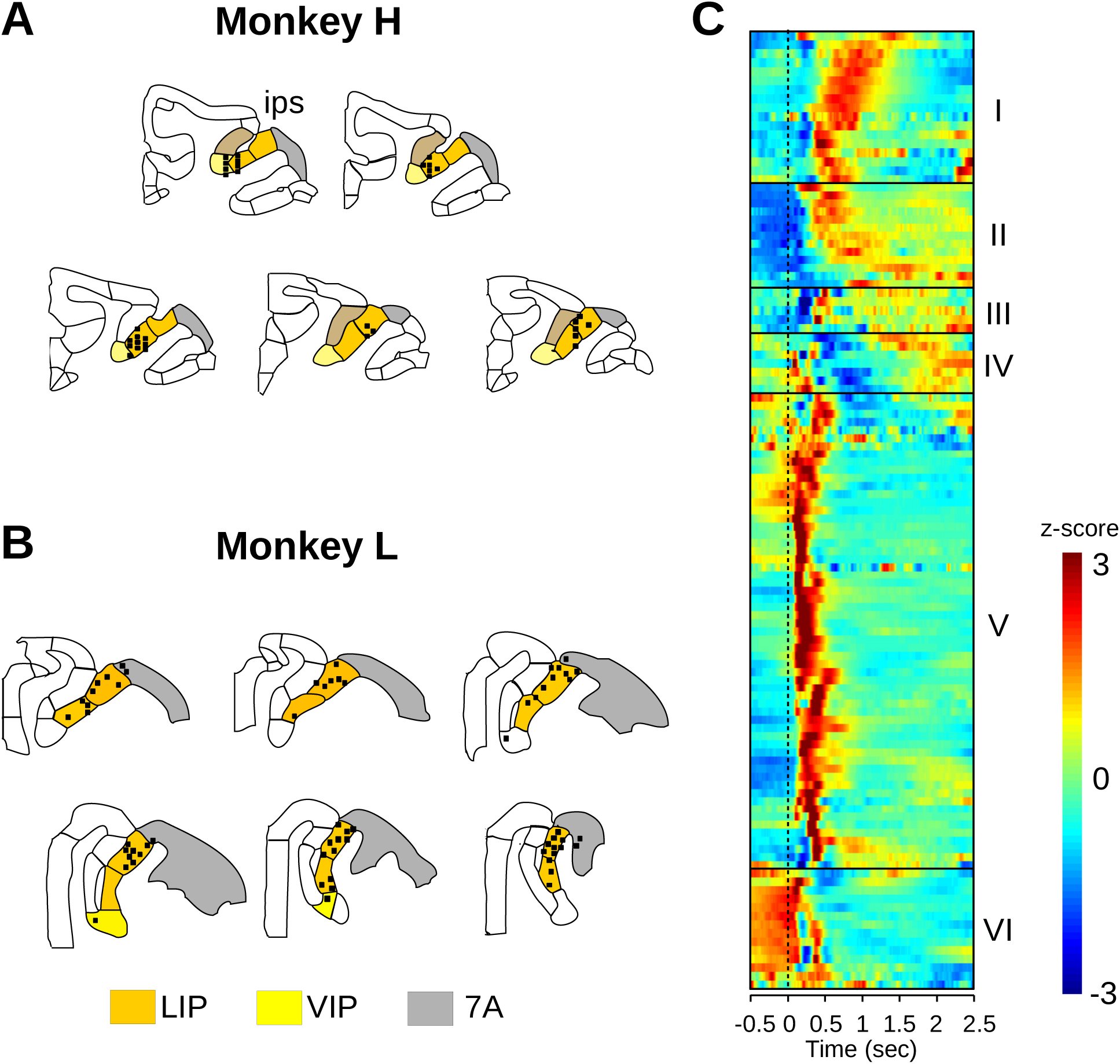
Reconstruction of recording sites and classification of neurons. **A**,**B.** Anatomical reconstructions of recording sites (*n* = 91) were based on stereotaxic MR images, chamber implantation and angles, and matched to the macaque rhesus anatomical templates (Reveley et al., 2017). The templates were adapted to overlap over the MRI images plus the recording sites, which ranged from 6mm posterior to 6mm anterior to the interaural plane. Amber regions are area LIP, yellow is VIP, and gray is area 7a. **C.** Six families of response types were found based on hierarchical clustering of mean firing rate vs. time, across all conditions. The neurons were optimally ordered using correlation similarity.

### Statistical Analysis

Analysis of behavioral and neuronal data consisted mainly of the following techniques: bootstrapped permutation testing (BPT), kernel density estimation (KDE) and Gaussian process regression (GPR). BPT was used to determine whether stimulus labels were informative with respect to overall neuronal activity. KDE was used for session-level averages of specific epochs within each trial, in which fairly large numbers of observations could be mustered to obtain an optimized nonparametric estimate. GPR was used to make inferences about processes that unfolded trial by trial, in order to interpolate across gaps.

Each pair of stimuli was classified according to two metrics: symbolic distance (a measure of relative position, calculated by taking the difference between the stimulus ranks) and joint rank (a measure of absolute position, calculated by taking the sum of stimulus ranks). Because these metrics are strictly independent of one another, they form a two-dimensional continuum across which performance can be analyzed (Jensen et al., 2017).

#### z*-transform of neural activity*

To construct a common metric across recording sessions with different activity levels, the firing rate on each trial (*FR*_trial_) was converted to a *z*-score using the mean of the firing rate (*µ*_*FR*_) and standard deviation (*σ*_*FR*_) over the entire session:

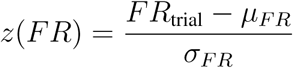

#### Bootstrap testing

Bootstrapping was used to estimate distributions of sample means. *N* samples were randomly drawn with replacement from the original set of *N* observations. The mean of the resampled observations was calculated. This procedure was repeated typically 10,000 times to generate the bootstrap distribution.

#### Random permutation testing

When examining the relationship between neural activity and explanatory variables such as ordinal position, it is useful to ask whether the observed relationship could arise by chance given the observed neuronal activity statistics. This was done by randomly permuting (“shuffling”) the explanatory variable labels among the trials. Each label occurred as many times in the shuffled data as in the original data. This procedure estimates the likelihood that apparent “tuning” for a particular explanatory variable could arise by chance. Random permutation testing could be combined with bootstrapping to obtain a distribution of apparent tuning functions.

#### Kernel density estimation

Eye movement characteristics and average session rates of neural activity both used KD estimates. For eye movements (position and velocity), we used a Gaussian kernel with Silverman’s optimal bandwidth (Silverman, 1986). For spike trains, we used the method described by Shimazaki and Shinomoto (2010).

#### Gaussian process regression

To evaluate how estimated performance changed as a function of learning, we used Gaussian process regression (Rasmussen and Williams, 2006). GPR is a highly flexible non-linear estimation technique that is well-suited to time series analysis. Although not widely used in neurophysiology, GPR is a well-established procedure (Durbin, 1985) that has seen extensive application in other domains. It has been called a categorical analysis with an infinite number of categories, arrayed along a continuum (McElreath, 2016). One continuum of interest is time: Response accuracy for each pair changes over time. Orthogonal to time are the continua of symbolic distance and joint rank, which are also expected to influence response accuracy. GP regression is performed by estimating the extent to which every observation covaries with every other, given some prior metric for comparing the distance between observations along the continua of interest. Each observation influences the estimate for other ‘nearby’ observations (e.g. that occur at similar times or have similar symbolic distance) more than observations that are distant (Lucas et al., 2015). Although such an analysis is not possible using classical methods (because of irreducible uncertainty about how distance and similarity interact to give rise to the data), Bayesian methods make GPR feasible by imposing a strong prior belief that pairs at similar times and with similar symbolic distance and joint rank should display similar patterns of neural activity (Gelman et al., 2014). GPR depends on relatively few assumptions, instead allowing the data to determine the form taken by the time series estimate. The chief constraint is that Gaussian processes are presumed to be smooth (i.e. differentiable without discontinuity). Beyond this constraint, one can imagine the model estimate as the posterior distribution of the relative density of all possible smooth curves, conditioned on the data and the informative prior. Although a full Gaussian process model can be computationally prohibitive to fit (requiring runtime *O*(*n*^3^) for *n* observations), we took advantage of the “expectation propagation” approximation (Tolvanen et al., 2014) implemented in the GPstuff toolbox (Vanhatalo et al., 2013) to accelerate estimation.

## RESULTS

Two subjects (H and L) completed 141 behavioral sessions (NHP H: 74, NHP L: 67). The average number of trials per session was 711 (H) and 890 (L). In each session the subject learned a novel TI list; one session = one TI list. A total of 117 neurons were recorded (H: 64, L: 53) during these sessions. Some neurons were recorded with more than one TI list, and some sessions contained data from more than one neuron, separated using the spike sorting methods described above. The result was a total of 142 recordings (H: 62, L: 80), where each recording contains data from one neuron with a novel 7-item list.

### Behavioral Performance

A mental representation of an ordered list could be implemented as a direct representation of ordinal position, or by related variables such as symbolic distance and joint rank. Performance accuracy and response time were analyzed to determine the behavioral influence of these variables, as well as variables related to reinforcement. A total of 141 sessions were analyzed (H: 74, L: 67). Each session started with a novel set of pictures that were arranged into a list by assigning a rank order to each picture. The ranking was initially unknown to the subject. Figure 3 illustrates the development of both the probability of choosing a given stimulus and the probability of a correct response over the first 500 choice trials, estimated using GPR. The influence of item rank on choice probability emerged over the first 300 trials (Fig. 3A). Learning curves (Fig. 3B) showed a monotonic increase in response accuracy (Fig. 3B black line) from a baseline of 0.5 (chance) up to an average around 0.75. The shaded regions represent the 80% and 99% credible intervals around the estimates of the mean.

**Figure 3.**
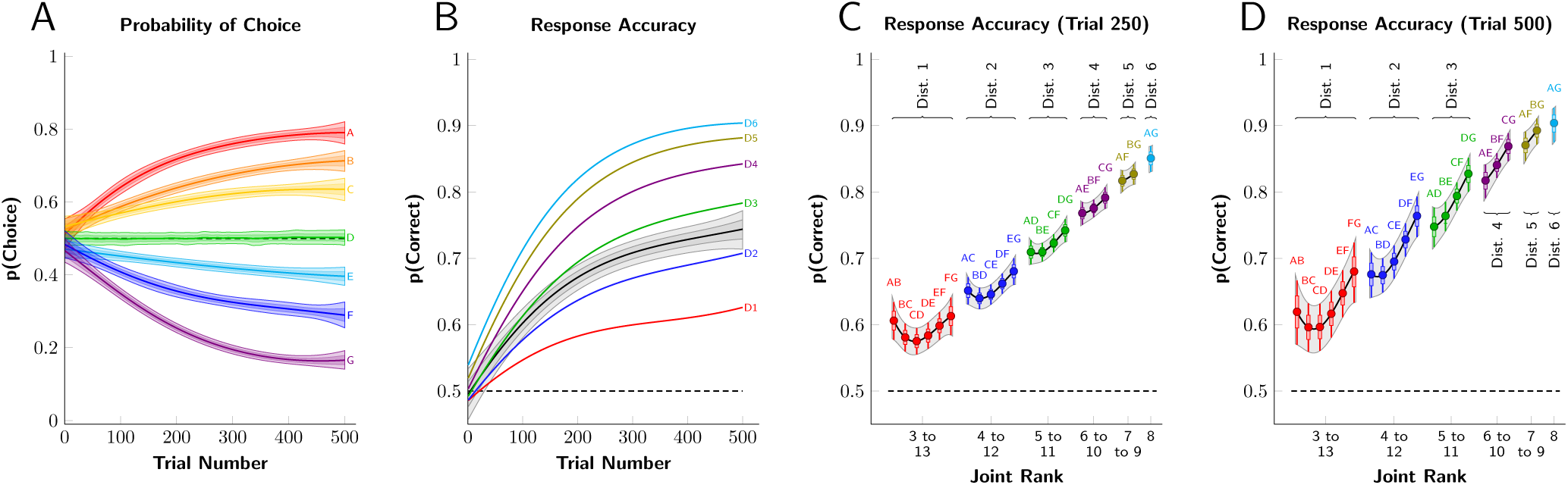
Response accuracy (proportion correct) was modeled as a Gaussian Process with a logit link, using trial number, symbolic distance, and joint rank as predictive dimensions. **A.** Probability of choosing each item during the first 500 trials. The 80% and 99% credible intervals for the mean are plotted as the dark and light shaded areas, respectively. **B.** Mean accuracy over the first 500 trials (black line), as well as accuracy sorted by symbolic distance (colored lines). **C.** Estimated accuracy for each pair at trial 250 sorted by joint rank and symbolic distance. Thick whiskers correspond to the 80% credible interval, and thin whiskers (as well as shading) correspond to the 99% credible interval. **D.** Estimated accuracy at trial 500.

Symbolic distance had a strong influence on performance accuracy (Fig. 3B,C), emerging with a time course similar to that of overall performance. Accuracy was lowest for adjacent pairs, somewhat higher for distance 2 pairs, and so on (Fig. 3C). Across sessions, the correlation (Pearson’s *r*) of overall performance (session average) with symbolic distance was significant for both subjects (H: *r* = 0.62, *p* < 0.0001, *N* = 400; L: *r* = 0.61, *p* < 0.0001, *N* = 402) with moderate effect sizes (H: *r*^2^ = 0.38; L: *r*^2^ = 0.37).

The correlation of overall performance with joint rank was significant for only one subject and effect sizes were very small for both animals (H: *r* = 0.04, *p* = 0.32, *N* = 726; L: *r* = 0.15, *p* < 0.0001, *N* = 737). Performance tended to be better for pairs that included the first and last list items, a “terminal item effect” (Fig. 3C & 3D).

Reaction times were remarkably short. Even though subjects were allowed up to 1.5 sec after stimulus onset to respond by making a saccadic eye movement to the target or distractor, choice reaction times (RT, aka saccade latency) were 232 +/- 31 msec (mean +/- s.d.) and remained in a narrow range throughout each session. Choice reaction time varied inversely with symbolic distance (ANOVA *p* < 0.0001, *N* = 87, 296), but the range was only 7 msec between the longest and shortest mean RTs (distance = 1, mean RT = 233 msec; distance = 6, mean RT = 226 msec).

Overall, these results showed that subjects rapidly acquired a representation of the list order, consistently expressing a preference for items of lower rank after only 250 to 300 trials of experience with an otherwise never-before-seen set of stimuli.

### Electrophysiology

We recorded from every neuron that we were able to isolate. We categorized neurons using a hierarchical clustering algorithm based on similarity of firing rate dynamics. Firing rates were first averaged across trials using KD estimation (Shimazaki and Shinomoto, 2010), to obtain an overall temporal response profile for each neuron. These averages were then *z*-standardized, and sorted into clusters based on the similarity of their temporal profiles. Categories were identified by building a dendrogram of correlations using agglomerative hierarchical cluster analysis (Hastie and Tibshirani, 2010), with *r* = 0.7 as the cutoff for cluster membership. This resulted in 6 clusters, depicted in Figure 2C. All classes of neuron were included in subsequent analyses.

Cluster V, comprising visual neurons, contained the most recordings. An example recording session for one visual neuron is shown in Figure 4. The cell had a transient, short latency (50 msec) response to the appearance of a visual stimulus in its receptive field (Fig 4A). The neuronal response started much earlier than the behavioral response latency (150 msec for single targets, 233 msec for choice targets). Spatial selectivity (Fig. 4B) was assessed on single target trials, showing a 2:1 difference between responses to stimuli inside (Fig. 4B blue) vs. outside (Fig. 4B magenta) the receptive field. On choice trials, selectivity was assessed by sorting according to saccade direction (Fig. 4B, black, grey). This cell did not differentiate between saccades toward or away from the RF, and can thus be said to have visual spatial selectivity, but not movement-related selectivity. In Figure 4C, firing rate has been converted to *z*-score and plotted as a function of joint rank (the sum of the ranks of the two stimuli presented in each trial) for trials with two stimuli (choice trials). Responses on choice trials are also plotted as a function of symbolic distance (difference in rank of the two stimuli, Fig. 4D) Figure 4E plots response vs. ordinal position (rank) for the stimulus inside the receptive field. During single target presentations (Fig 4E, black), variations in the response to different images can only be due to the visual properties of the stimuli, as the NHP had yet to learn the rank of each stimulus. A *z*-score of zero implies that the response was equal to the average response across all conditions. This cell shows effects that were characteristic of the population: strong spatial selectivity for single targets, response averaging for two stimuli, weak selectivity for ordinal position, and modulation by symbolic distance and joint rank. These features of the data are elaborated below.

**Figure 4.**
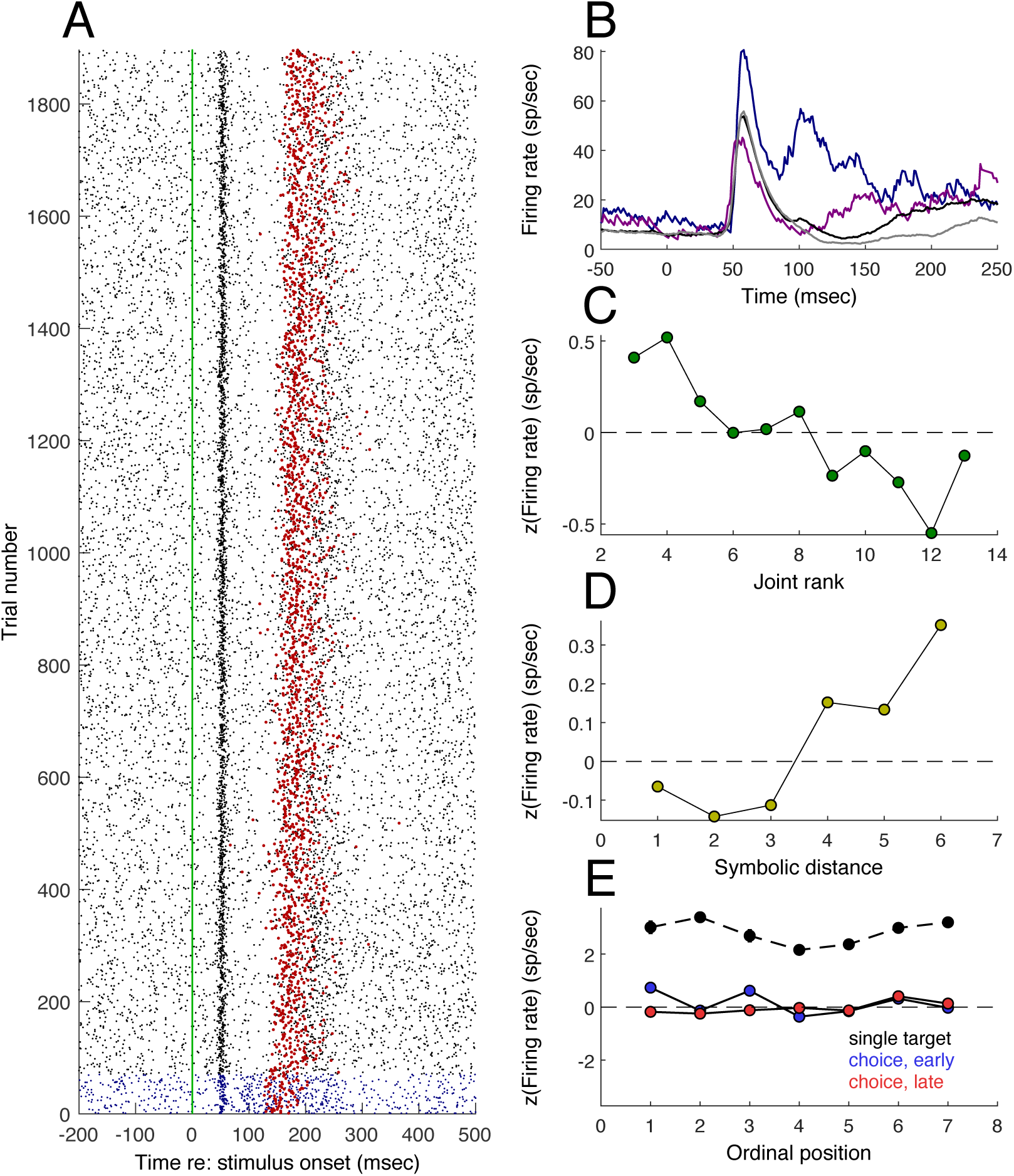
Example of a recording session for one cell. **A.** Spike raster aligned on stimulus onset (vertical blue line). Each blue dot (for single target trials) or black dot (for choice trials) represents an action potential. Each row is one trial. Vertical green line is stimulus onset. Red dots are saccade latencies. **B.** Smoothed average firing rate vs. time. Blue line is for a single target in the receptive field, purple for a single target outside the receptive field. Black/gray corresponds to choice trials (two targets) where the saccade was toward (black) or away (gray) from the receptive field. **C.** *z*-scored firing rate vs. ordinal position. **D.** *z*-scored firing rate vs. symbolic distance. **E.** *z*-scored firing rate vs. joint rank.

#### Encoding of joint rank and symbolic distance

For each recording, *z*-scores of the average firing rate in the interval starting 50 msec after stimulus onset and ending with saccade onset were sorted by joint rank. An example recording with choice trials sorted by joint rank is shown in Fig. 5A. Figure 5B shows the firing for each recording, sorted by the joint rank that was associated with the strongest response. Figure 5C shows the decoding of joint rank by an optimal linear estimator (OLE, Salinas and Abbott, 1994) using the neuronal ensemble activity. The OLE was run 1,000 times and the variability of estimates was obtained with a bootstrap procedure. The same analyses were done for symbolic distance (Fig. 5D,E,F). Roughly half the recordings had the greatest activity for one of the two largest symbolic distances (Fig. 5E). Across the population most neurons tended to have higher firing for larger symbolic distance, whereas the tuning for joint rank was more distributed. As was the case for the behavioral results (Fig. 3C,D), the neural correlates of joint rank and symbolic distance emerged during the first 250 learning trials and were stable for the duration of the session.

**Figure 5.**
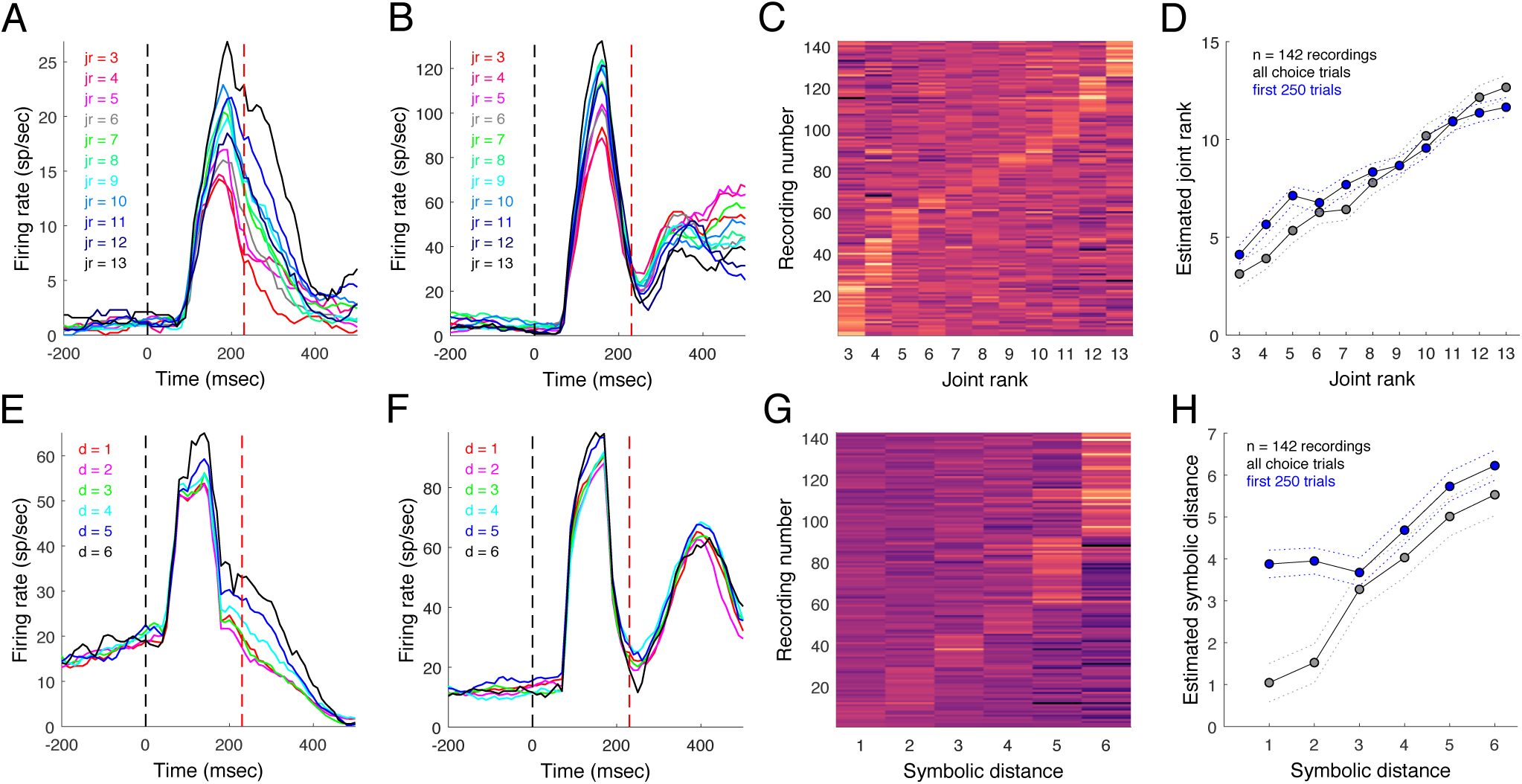
Joint rank and symbolic distance coding. **A.** Example post-stimulus time histogram for one recording with choice trials sorted by joint rank. **B.** Example neuron for second monkey. **C.** Recordings (*n* = 142) sorted by preferred joint rank based on *z*-scored firing rate. **D.** Estimated joint rank using an optimal linear estimator. Gray lines are 95% confidence intervals for all trials based on 10,000 bootstrap iterations. **E.** Example post-stimulus time histogram sorted by symbolic distance (different session than **A**). **F.** Example neuron for second monkey. **G.** Recordings sorted by preferred symbolic distance. **H.** OLE estimated symbolic distance with bootstrap estimates of standard error.

The effects of joint rank and symbolic distance were further quantified by computing the variance accounted for (VAC) by each variable. For each recording, firing rate was smoothed by applying Gaussian Process Regression (GPR) using two models; one with trials, time, and Joint Rank (JR), and another with trials, time, and symbolic distance (SD). Because GPR is flexible and non-linear, a measure of the VAC provided a way of assessing the *degree* of tuning a cell displayed as a function of that variable, without specifying a particular *form* that tuning had to take. The estimation was done within 4 epochs: baseline (250 ms before stimulus onset), visual (between 5 ms to 150 ms after visual presentation), presaccadic (100 msec before saccade initiation) and postsaccadic (500 ms after saccade execution) period. Each period was divided into intervals of 10 msec to create a spike counts over time as a prior to the GPR the posterior estimate of firing rate (mean and standard deviation) was a continuous function of time, trial, JR and SD. The estimated firing rate and uncertainty were then averaged within the previously specified time windows. To estimate the amount of variance associated with JR or SD across the session for visual, presaccadic or postsaccadic epochs, we subtracted the baseline, and then evaluated the variance accounted for (VAC) using partial eta-squared 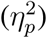, with *F* -values and degrees of freedom from the Welch test for populations with unequal variance. Each 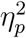 was sorted by cell class (see Fig. 2C) and in order of increasing VAC magnitude (Fig. 6 A-F). The density of each 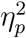 was estimated by bootstrap. VAC values for JR were generally larger than for SD. VAC was similar across clusters, suggesting that JR and SD are represented across all cell classes.

**Figure 6.**
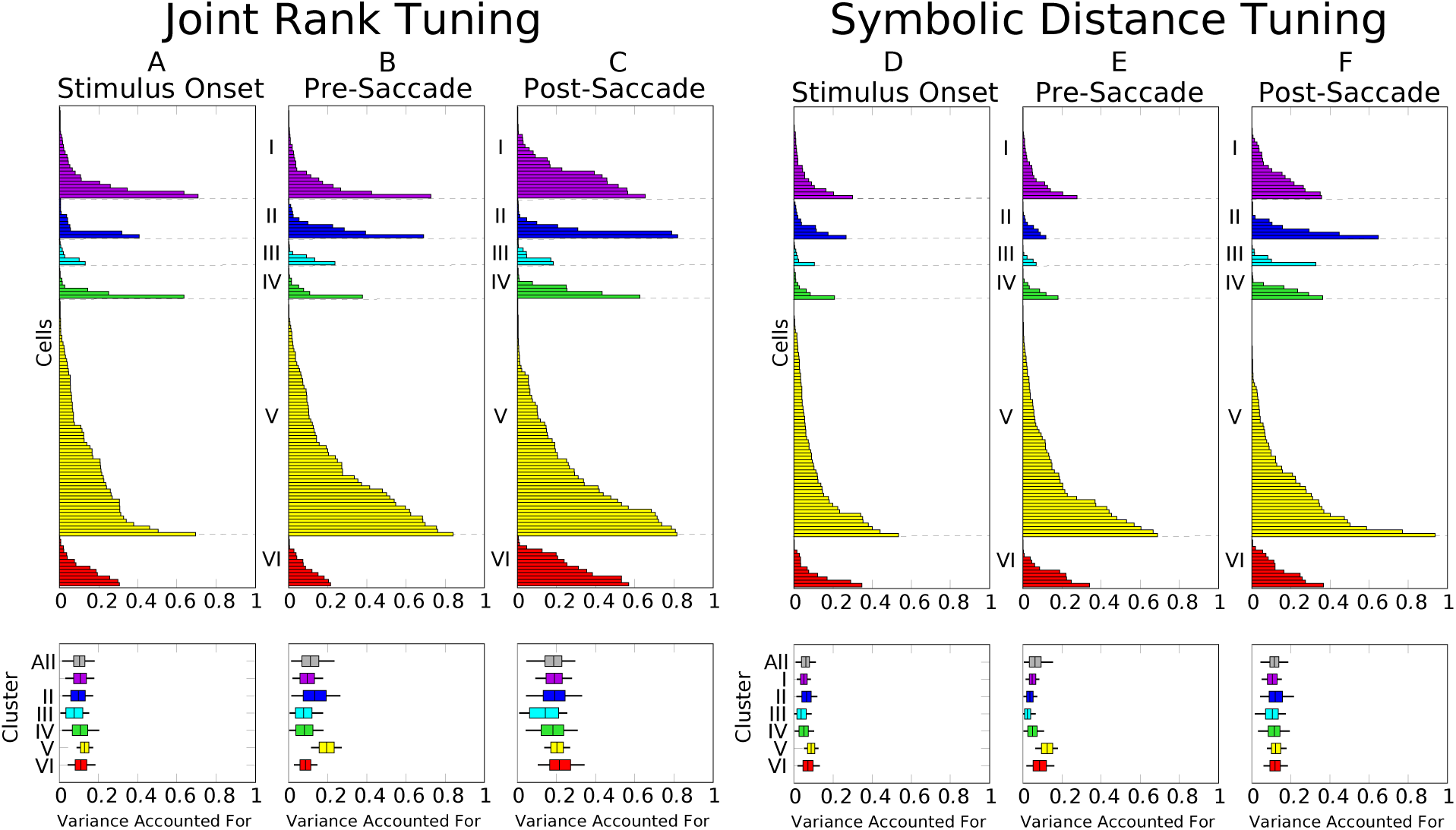
Variance accounted for by joint rank and symbolic distance, sorted by neuron class and by trial epoch. Box-and-whisker diagrams depict mean VAC for each cluster, with boxes corresponding to 80% credible intervals and whiskers corresponding to 95% credible intervals.

Joint rank and symbolic distance are uncorrelated by design. Therefore, it is possible that individual neurons could be modulated by either or both variables. In Fig. 7, the VAC for JR is plotted against SD on a cell-by-cell basis for three trial epochs. Both variables are more strongly represented around the time of the choice saccade than around the time of stimulus onset, as suggested by Fig 5A and 5D. The variance accounted for by both factors tends be correlated across the population of recordings. Hence, neurons representing JR and SD are not distinctly separate subpopulations. However, the vast majority of neurons have greater VAC for JR than for SD across all trial epochs.

**Figure 7.**
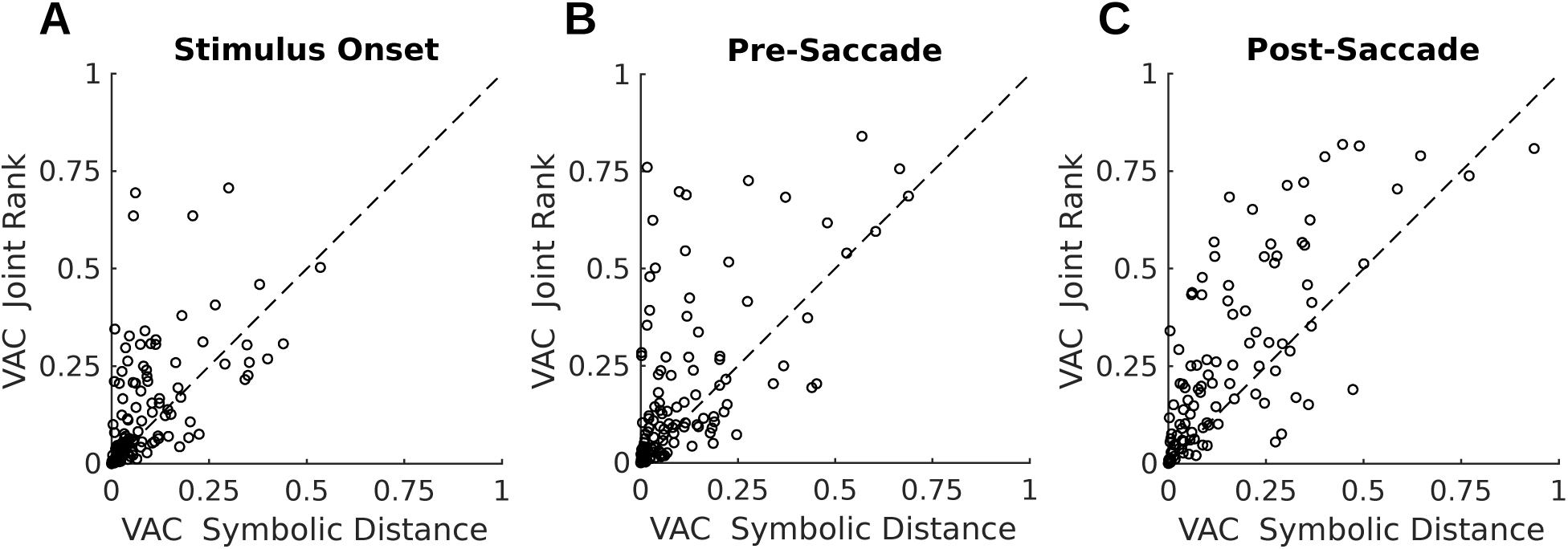
Variance accounted for (VAC) by Joint rank vs. symbolic distance. *n*=142 recordings with 7-items lists. **A.** VAC in firing rate activity aligned to stimulus onset. **B.** VAC in activity prior to saccade onset. **C.** VAC in activity prior to saccade onset.

#### Encoding of stimulus identity or rank

Sereno and Maunsell (1998) reported visual feature selectivity in monkey posterior parietal cortex using stimuli that controlled for luminance cues. In the current study, the stimuli were not controlled for luminance, chromaticity, contrast or spatial frequency content. They were simply random colorful images with content that was easily recognized and discriminated. Nevertheless, it is possible to ask if neuronal responses to the different stimuli varied reliably. The fact that the stimuli varied along multiple visual dimensions increases the expectation that such neural response variability should be found. For this analysis, *n* = 142 sessions were used. Spikes were counted within a time window 50-150 msec after stimulus onset on each trial, converted to spikes per second, and then *z*-scored.

Trials in which only a single stimulus appeared (i.e. the first 5 blocks of each session) were sorted according to the identity of that stimulus. Figures 8A shows the values of *z*(FR) sorted by stimulus and averaged over all recording sessions. The stimulus that evoked the strongest response in each session was assigned to position 1, and the other stimuli were assigned to positions 2-7, maintaining their relative rank order. Some degree of apparent stimulus preference can arise by chance, given finite sampling of variable neural responses. To control for this, the preceding analysis was repeated with the stimulus identities randomly shuffled among trials. There was no significant difference between the shuffled and unshuffled responses (bootstrapped *p* = 0.58 for stimuli inside the RF; outside RF bootstrapped *p* = 0.70; post-hoc tests for each item were n.s.). Response differences across stimuli thus appear to be accounted for by random variability, and there is no evidence that neuronal responses reliably encoded stimulus identity during the initial phase of the task when the stimuli were presented individually.

**Figure 8.**
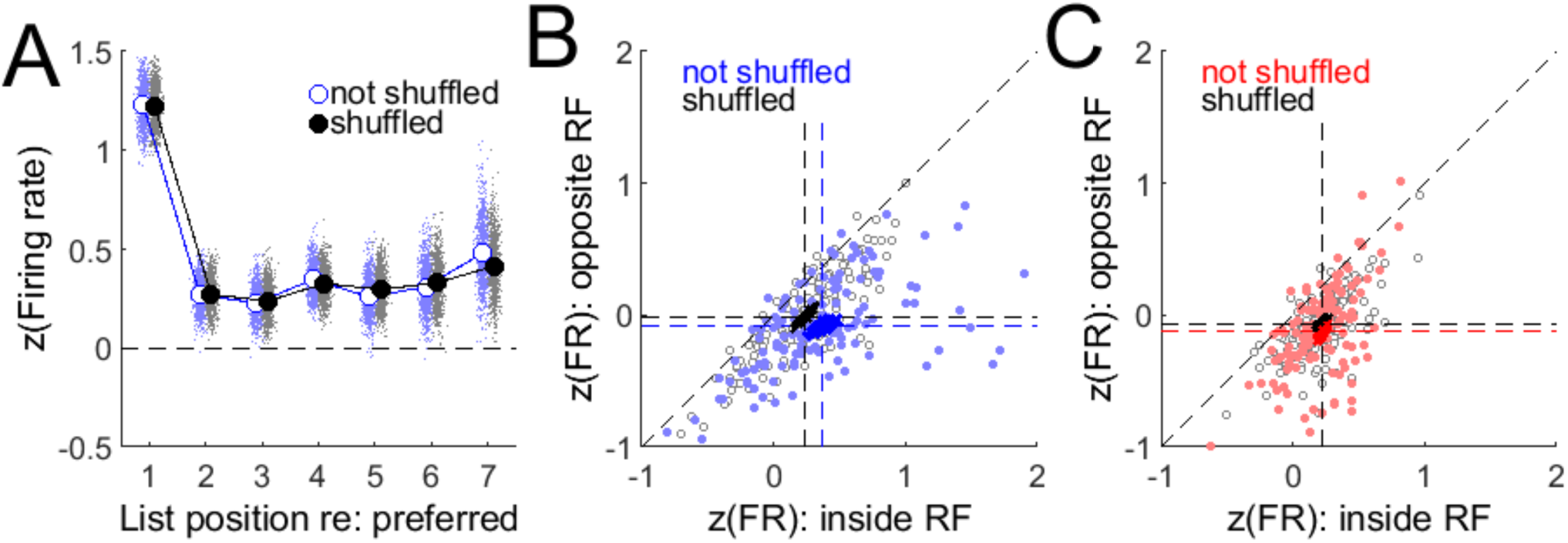
Encoding of stimulus identity. *n*=142 recordings with 7-items lists. **A.** Trials with a single stimulus in RF. Mean *z*-scored firing rate during visual epoch for original stimulus labels (blue, not shuffled) and shuffled labels (black). Light blue and grey points are estimates of the means from 10,000 bootstrap iterations. **B.** *z*-scored firing rate during visual epoch of first 250 choice trials with best stimulus inside RF. Light blue dots are responses with original labels, grey dots are responses with shuffled labels. Dark blue and black dots are estimates of the mean from 10,000 bootstrap iterations. **C.** Same as B, but for choice trials after trial 250.

After the first 5 blocks of each recording session, images were presented in pairs, one image inside of the receptive field and the other opposite to the receptive field. As shown above (Fig. 3), TI learning was most rapid during the first 250 trials of paired presentations. During these trials, there was a single best stimulus for each neuron that evoked a stronger response compared to the “best” stimulus when stimulus identity was shuffled across trials (Fig 8B, bootstrapped difference *p* < 0.001, *n* = 142). This effect was present during learning trials (i.e., the first 250 paired presentations, Fig 8B), but was not present during later trials, when performance reached a plateau (Fig. 8C; bootstrapped *p* = 0.8; *n* = 142). Hence, during TI learning, visual responses transiently but reliably encoded stimulus rank.

#### Effects of performance

Reward-related modulation of neural activity in macaque parietal cortex was described by Platt and Glimcher (1999). It has also been reported that neuronal activity in parietal cortex is correlated with performance accuracy (Zhang et al., 2014). Given current evidence for the influence of serial order and related quantities on both behavior and parietal neuronal activity, it is important to ask whether these findings could be related to reward associations. In the TI task, all correctly completed trials were rewarded, thus reward probability was equal to percent correct. To test for effects of performance or reward, we compared the difference in activity (spike counts) between correct and incorrect trials for each recording. Three trial epochs were examined: background (200 msec prior to stimulus onset), visual (50-150 msec after stimulus onset) and presaccadic (100 msec prior to onset of choice saccade). Each of 142 recordings was subjected to a Wilcoxon test with a criterion of *α* = 0.05. Effect sizes were measured using *η*^2^. Few recordings showed significant effects of performance in any trial epoch and the effect sizes were tiny (background epoch: *N* = 11 (8%) significant recordings, mean *η*^2^ = 0.003; visual *n* = 13 (9%), *η*^2^ = 0.003; presaccadic *n* = 25 (18%), *η*^2^ = 0.04), suggesting negligible association between neural activity and performance outcome or reward probability in the current data set.

These group differences were supported by Kruskal-Wallis tests run for each recording and explanatory variable (target location, saccade direction, outcome) with alpha = 0.05. Approximately 43% of cells (61/142) significantly different in the variance accounted for by target location, with an average effect size (generalized *η*^2^) of 0.34. Additionally, 35% (54/142) differed significantly as a function of saccade direction (avg. effect size = 0.08), and 20% (29/142) differed significantly as a function of outcome (avg. effect size = 0.02).

#### Effects of prior reward history

It is possible that subjects used a model-free reinforcement learning (RL) strategy to perform the task. A hallmark of model-free RL is that choices that are rewarded are more likely to be repeated than those that are not rewarded. We investigated this by first looking at the pattern of spatial choices; if the subject was rewarded for choosing the stimulus in a particular location, would they be more or less likely to choose the same location on the next trial regardless of whether it was the correct response? There was evidence in favor of a spatial win-stay, lose-shift bias; rewarded response directions were repeated 56% of the time, significantly different from the expectation of 50% (*t*-test *p* < 0.0001, *n* = 142, Cohen’s *d* = 0.68). Unrewarded trials were followed by responses to the same location only 48% of the time (*t*-test *p* < 0.002, *n* = 142, Cohen’s *d* = 0.27). This result generally agrees with a previous study of reward-based decision-making (Kubanek and Snyder, 2017), which found that monkeys were more likely to change their response after receiving a relatively small reward. Reaction time (choice saccade latency) did not vary depending on the outcome of the previous trial (mean RT difference, after previous reward - after no previous reward, = −0.9 ms, SD = 6.3 ms, paired *t*-test *p* = 0.1, *n* = 142).

It should be noted that a spatial choice bias is deleterious to overall performance, as the target location was randomized and provided no information about the correct response. The presence of a spatial bias does not rule out a choice bias that might facilitate learning of serial order. This possibility was tested by asking whether reward history biased the choice probability for each of the 7 pictorial stimuli conditioned on whether or not the previous choice of that stimulus had been rewarded. This analysis was performed for non-terminal list items (B thru F), for which the expected choice probability is 0.5, regardless of performance accuracy. Some conditions were excluded if the choice probability could not be calculated (i.e. the denominator was 0, which happened in 11 of 661 cases). The average probability of choosing a particular stimulus following a rewarded choice of that stimulus was 0.51 (SD = 0.17), compared to 0.51 (SD = 0.18) following unrewarded choices. The difference was not significant (*t*-test paired by session and item, *p* = 0.99, *n* = 650). Similarly, the average latency of choice saccades varied by less than 1ms between previously rewarded and unrewarded conditions (paired *t*-test *p* = 0.34, *n* = 661). The results were the same whether all choice trials in each session were included, or only the first 250 trials when learning of serial order was most rapid. These results provide no support for the idea that subjects used a model-free reinforcement learning strategy to learn or perform the task. If anything, rewarded choices were slightly less likely to be repeated than unrewarded choices.

The behavioral result that rewarded spatial choices were likely to be repeated (although rewarded list item choices were not) raised the possibility that reward history might affect neuronal responses. In a task with varying reward magnitudes, Kubanek and Snyder (2017) found that LIP neurons fired more vigorously following trials that ended with relatively small rewards. In the current data (142 recording sessions), there was little evidence that reward outcome affected neural activity on the next trial. For choice saccades toward the receptive field, firing rate during the period between stimulus onset and choice saccade was not significantly elevated on the trial following a rewarded trial compared to an unrewarded trial (reward - no reward, mean firing rate difference = 0.07 sp/sec, *p* = 0.75, *n* = 142). For saccades away from the receptive field, firing rate was non-significantly reduced (reward - no reward, mean firing rate difference = −0.19 sp/sec, *p* = 0.39, *n* = 142). These effects were negligible and suggest that, in the current data, reward history was not a significant modulator of neural activity.

## DISCUSSION

The transitive inference (TI) paradigm tests subjects’ ability to learn the implied serial order of a set of pictorial stimuli without any explicit spatial or temporal cues. This is accomplished by presenting pairs of images drawn from a rank-ordered list of images and rewarding subjects for choosing the item that has the lower rank. Much work in humans and other animals has supported the idea that TI learning relies on a virtual workspace (reviewed by Jensen, 2017), but few studies have examined its physiological underpinnings in brain regions implicated in spatial cognition (Brunamonti et al., 2016). Here, we tested the virtual workspace hypothesis by recording neuronal activity in posterior parietal cortex while monkeys learned novel TI lists consisting of sets of 7 images that were assigned ranks learned by trial-and-error.

Each recording session started with a novel set of images. Subjects initially responded randomly, but were able to learn each list within about 250 trials. Prior research argues against the idea that monkeys memorize the response to each pair (Treichler et al., 2007) or rely solely on the experienced reward value of each stimulus (Jensen et al., 2019). Rather, they appear to acquire abstract knowledge of the list order, and of the rule that dictates choosing the lower ranked item in each pair. In the current study, behavior was more consistent with representation and rule-based learning than with model-free reinforcement learning. Neuronal activity in parietal cortex displayed strong visual and spatially selective, but also showed significant influences of cognitive variables related to list order such as item rank, symbolic distance, and joint rank. These results support the idea that parietal cortex is involved in representing abstract serial order.

The strength of the observed joint rank effects was not expected at the outset of the study, because joint rank is much less predictive of response accuracy than symbolic distance. The stimulus pairs AG, BF, and CE all have the same joint rank of 8, for example, but differ considerably in their levels of response accuracy (Fig. 3). Despite not providing an obvious benefit to the downstream behavior, however, it may be that joint rank is encoded as part of a wider scheme for interpreting complex scenes that contain multiple stimuli. Just as symbolic distance and joint rank can be calculated arithmetically from the ranks of the original stimuli (SD=B-A, JR=B+A), the ranks of the stimuli can be decomposed from an encoding of symbolic distance and joint rank (B=(JR+SD)/2, A=(JR-SD)/2). Cells that encode these additive and subtractive quantities from across the visual field could be processed downstream to represent individual stimulus identities, and parietal cortex may not be the locus of that decomposition.

Joint rank may also be related to an approximate number system in monkeys (Brannon and Merritt, 2011; Cantlon, 2012). Neurons related to the representation of visual quantity have been described in the parietal cortex of macaques (Nieder et al., 2002; Nieder, 2013; Roitman et al., 2007, 2012). This representation may support simple arithmetic operations that are more fully developed in humans (Fehr et al., 2007; Rosenberg-Lee et al., 2011; Arsalidou et al., 2018).

We tested the contribution of model-free learning mechanisms by performing a sequential trial analysis to determine if received rewards biased subjects’ behavior on subsequent trials. Reward history was found to introduce a spatial bias, but did not bias responses in favor of choosing a previously rewarded list item. There was no evidence of neuronal responses being modulated reward history, as reported by others (Kubanek and Snyder, 2017). Variations in neuronal response were only weakly correlated with trial outcome (Zhang et al., 2014). These results are in accord with a sizable literature showing that experienced reward value doesn’t account for transitive inference or serial learning in multiple primate species (Terrace, 2012; Jensen et al., 2019).

There was no evidence that item identity based on shape, color, or other visual features was encoded when single stimuli were presented during the initial trials of each session (prior to TI learning), in contrast with previous work showing shape selectivity in LIP (Sereno and Maunsell, 1998). This was despite substantial item-wise variation in responses, possibly driven by low-level visual properties of the stimuli that were not systematically controlled. However, this variation did not survive a permutation test, which is a stringent control for reliable stimulus encoding.

During the first 250 trials of TI learning, neural activity reliably distinguished among list items. Because the neurons showed no inherent visual selectivity prior to learning, it is likely that this signal was related to the learned list positions. The effect of list position disappeared after the initial learning, as accuracy reached a plateau. Thus, there appears to be a transient representation of item list position in parietal cortex during learning. The lack of selectivity during single stimulus presentations argues against the possibility that this representation was driven by visual features of the stimuli.

The representation of variables that arise from the comparison of spatially remote stimuli is contrary to conventional thinking about visual or visual-movement neurons with spatially localized receptive or movement fields. However, such responses are not unprecedented in the dorsal visual pathway of macaques. Zaksas and Pasternak (2005) found effects of remote stimuli in visual area MT when monkeys were required to compare motion stimuli in the contra- and ipsilateral visual fields. Moreover, models of spatial remapping around eye movements strongly suggest that individual LIP neurons receive information from across the entire visual field (Quaia et al., 1998).

Overall, our results support an emerging view of parietal cortex in which: 1) Spatial and non-spatial information is represented by the same population of neurons. 2) Non-spatial information can encode abstract quantities related to implied serial order, which obeys the law of transitivity. Finally, 3) information is integrated across visual space to compute cognitive variables based on the comparison of spatially remote stimuli. All of these properties add to growing evidence that parietal cortex likely plays a substantial role in the construction of a virtual workspace for both spatial and abstract cognitive tasks.

## Acknowledgments

We are grateful to Drew Altschul and Jessica Joiner for assistance during the early part of this project.

## Author Contributions

The study was designed by GJ, VPF, and HST. Data were collected by FM, YA, and BCK. Data were analyzed by GJ, VPF, and FM. The paper was written by VPF, GJ, BCK, and FM, with input from HST and YA. FM and GJ made equal contributions.

